# Linking comparative genomics and environmental distribution patterns of microbial populations through metagenomics

**DOI:** 10.1101/058750

**Authors:** Tom O. Delmont, A. Murat Eren

## Abstract

Combining well-established practices from comparative genomics and the emerging opportunities from assembly-based metagenomics can enhance the utility of increasing number of metagenome-assembled genomes (MAGs). Here we used protein clustering to characterize 48 MAGs and 10 cultivars based on their entire gene content, and linked this information to their environmental distribution patterns to better understand the microbial response to the 2010 Deepwater Horizon oil spill in the Gulf of Mexico coastline. Our results suggest that while most oil-associated bacterial populations originated from the ocean, a few actually emerged from the sand rare biosphere. These new findings suggest that there are considerable benefits to employ approaches from comparative genomics to study the whole content of newly identified genomes, and the investigation of emerging patterns in the environmental context can augment the efficacy of assembly-based metagenomic surveys.

---

During the last two decades, the genomic content of more than 40,000 microbial isolates have been characterized and used to study the microbial gene pool, adaptation, and evolution [1, 2, 3, 4, 5]. Although cultivation-based methods have paved the way for the emergence of powerful comparative genomics approaches, comprehensive understandings of microbial distribution patterns and niche boundaries remained hard to achieve due to well-understood limitations of cultivation.

A complimentary solution emerges from assembly-based metagenomics, where both the genomic content, and the relative distribution of naturally occurring microbial populations can be recovered [6, 7]. To explore microbial systems using metagenome-assembled genomes (MAGs), researchers often rely on functional annotations, phylogenetically informative conserved gene families, or distribution patterns to identify metabolic potentials [8], evolutionary relatedness [9, 10], or co-occurrence of genomic collections [11]. Despite their efficacy, these approaches disregard the signal from genes that are not yet characterized, but may be critical for niche adaptation [12].

Characterizing often-novel MAGs by taking their entire gene content into consideration, in conjunction with their distribution recovered from metagenomic data could compliment current practices, and provide additional insights into the ecology of microbial ecosystems. Here we investigated MAGs and cultivar genomes associated with the 2010 Deepwater Horizon (DWH) oil spill to improve our understanding regarding the origin of oil-associated microbial populations using protein clusters (PCs), and their environmental distribution patterns.

The shotgun metagenomic dataset we studied contained 16 samples collected by Rodriguez-R et al. [13] from Pensacola Beach (Florida) (1) before the oil from DWH began to wash ashore, (2) during the oiling event, (3) and after the removal of oil from the beach. This dataset of 452 million reads was originally analyzed at the contig-level [13], and our re-analysis had resulted in the recovery of MAGs spanning various taxonomical groups [14]. Of those, 14 MAGs were mostly detected before or after the oiling event, while 34 MAGs and all ten cultivar genomes isolated from the same environment [15] were enriched only in oil-contaminated samples (Figure 1). The distribution patterns across environmental conditions revealed a strong link between the genomes enriched in oil-contaminated samples, and genes coding for urea metabolism [14]. Urea is a dissolved organic nitrogen compound used by marine microbes as a main source of nitrogen (Solomon et al., 2010), which prompted us to suggest that bacterial populations enriched in the Gulf of Mexico coastline during DWH originated from the ocean rather than emerging from the sand rare biosphere [14].

To revisit these findings, here we used 176,024 genes identified in the 48 MAGs and 10 cultivar genomes, and clustered them into 14,991 non-singleton protein clusters (PCs) (see Supplementary Methods). We then (1) clustered PCs based on their occurrence patterns across genomes, (2) organized genomes based on PCs they harbor, and (3) created a holistic display that conveys the distribution patterns of genomes in the environment (Figure 1). Five groups emerged from the organization of genomes based on the PCs they shared. Groups I-II, and IV were mostly enriched during oiling, and contained 32 MAGs and all 10 cultivar genomes. In contrast, most of the 16 MAGs in groups III and V were detected only before or after oiling.

26% of the 12,983 functions we identified distributed differentially between the five groups (p ¡ 0.05; Figure 1). We incorporated a subset of these functions into our display to highlight the biological relevance of groups (Figure 1). Among them, genes coding for urea metabolism were widespread in groups I-II, and IV but missing in groups III and V, providing a stronger partitioning of urea metabolism by PCs compared to distribution patterns alone (Figure 1). Groups I-II-IV were also enriched with genes coding for TRAP transporters (specialized in the uptake of organic acids), glutathione reductase and glutathione S-transpherase (oxidative stress and detoxification), revealing distinct nutrient acquisition and stress response strategies compared to groups III and V (Figure 1). Furthermore, genes coding for flagellar biosynthesis and twitching mobility were widespread in all groups except group III. Instead, group III was enriched in genes involved in gliding motility, providing a means to travel in environments with low water content ([16]).

**Figure 1.**
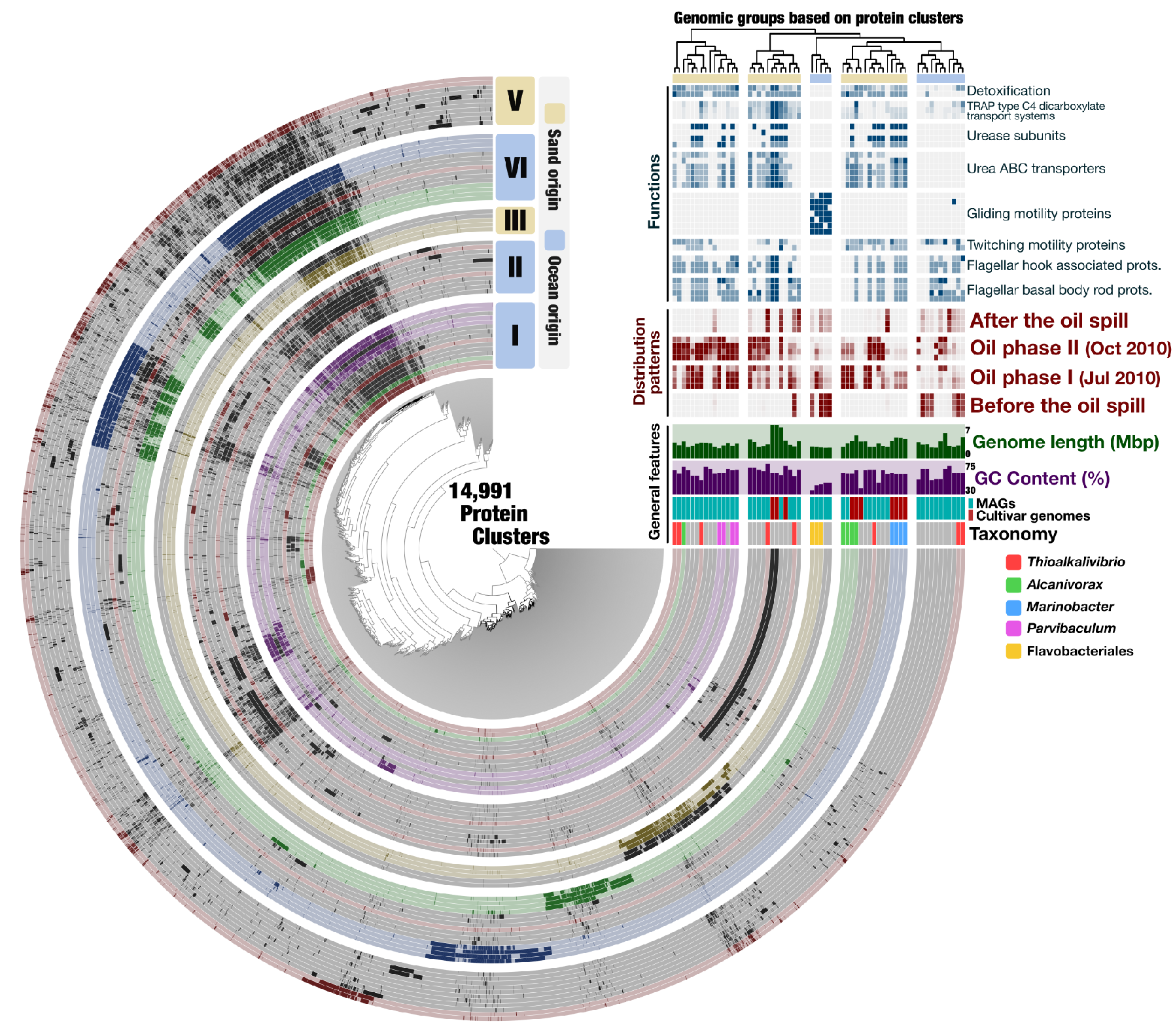
Clustering of 58 microbial genomes identified from the Gulf of Mexico coastline based on 14,991 protein clusters (PCs). Each radial layer represents a genome, and each bar in a layer represents the occurrence of a PC. The organization of genomes is defined by the shared PCs (the second tree on the right-top). In addition, general features, environmental distribution patterns, and a selection of differentially occurring functions are displayed for each genome. A high-resolution copy of this image is temporarily hosted here.

The overall coherence between the partitioning of genomes based on shared PCs, and their functional potential and distribution patterns, supports the hypothesis of a distinct origin for oil-associated microbes (Figure 1). This holistic strategy elucidates which bacterial populations likely originated from the ocean (groups I-II, and IV), and separates them from the native sand bacterial populations (groups III and V). Interestingly, groups associated with the sand ecosystem include genomes undetected before the oiling event, revealing a previously overlooked finding that some oil-associated populations emerged from the sand rare biosphere.

PCs also show improvement over taxonomy. For instance, eight *Thioalkalivibrio* genomes in the dataset occurred in four of the five genomic groups, reminding that co-existing microbial populations sharing the same taxonomical affiliation can strongly diverge functionally and may not necessarily originate from the same ecosystem. In another example, all three Flavobacteriales MAGs clustered in group III, yet one of them was oil-associated while the two other occurred mostly before the perturbation, suggesting an oil-triggered shift in the relative distribution of Flavobacteriales MAGs, possibly within the same niche.

In summary, combining comparative genomics, environmental distribution patterns, and functional potential of genomes in a single context allowed us to achieve a more detailed depiction of the ecology of microbes in an oil-challenged environment by enhancing the utility of MAGs. This approach revealed co-occurring microbial populations that originate from distinct ecosystems, and differentially occurring microbial populations that share the same one. Our findings also showed that although most of the oil-degrading microorganisms in Pensacola Beach originated from the marine environment, some others likely emerged from the sand rare biosphere. These new findings demonstrate the benefits of studying the whole content of newly identified MAGs, and investigating emerging patterns in the environmental context.

## Acknowledgements

We thank Julie Reveillaud for her comments on our manuscript. As two research parasites, we have the utmost respect and gratitude towards Rodriguez-R et al., Overholt et al., and others who studied the 2010 Deepwater Horizon oil spill and made their data publicly available.

## Methods

**Genomes and metagenomes.** The sand metagenomes were generated by Rodriguez-R et al. [13] and includes four samples collected before the oil began to wash ashore (May 2010), eight samples collected during the oiling event (four in July 2010, and four in October 2010), and four samples collected after removal of oil from the beach (June 2011). Raw metagenomic sequencing data for these 16 samples are publicly available under NCBI BioProject ID PRJNA260285. Data for ten cultivar genomes from Overholt et al. [15] are publicly available under NCBI BioProject ID PRJNA217943. Cultures were grown using oil as the sole source of carbon from samples collected from Pensacola Beach and other Florida beaches affected by the oil spill. FASTA files for 48 MAGs [14], ten cultivar genomes, and anvi’o files to reproduce Figure 1 are publicly available via https://dx.doi.org/10.6084/m9.figshare.3199459.

**Recovering the distribution patterns of genomes and generating protein clusters.** After noise filtering (see [14] for details), short reads from each metagenomic sample were mapped back to cultivar genomes and MAGs using CLC Genomics Workbench (version 6) (http://www.clcbio.com) by requiring a minimum of 97% sequence identity over 100% of the read length. Anvi’o reported the mean coverage and portion coverage of each genome in all metagenomic datasets. To generate protein clusters we used ITEP [17] (with inflation and maxbit parameters set to 2.0 and 0.1) to profile GenBank files for the 10 cultivar genomes and 48 MAGs using a server running Linux CentOS version 6.4. Figure 1 reports the occurrence of protein clusters across genomes.

**Taxonomic and functional annotation.** We used the RAST platform [18] to infer the taxonomy and functionality of genomes, and to acquire GenBank files that contain the location and the translated protein sequence for open reading frames identified in each genome. We used STAMP [19] to perform ANOVA test to determine differentially occurring functions between groups of genomes as identified by protein clusters.

**Generating the anvi’o display and network visualization**. We used anvi’o v1.2.2 (available from http://github.com/meren/anvio) to analyze protein clusters and integrate the environmental data. We transformed the ITEP tab-delimited output for protein clusters into an anvi’o compatible format using the script `anvi-script-itep-to-data-txt`. We then created a hierarchical clustering of protein clusters based on their distribution across the 58 genomes using the script `anvi-matrix-to-newick`, which employed Euclidean distance as distance measure. We used `anvi-gen-samples-info-database` to create an additional anvi’o database to display basic features of genomes (i.e., GC-content, or genome length), as well as presence of select functions, and environmental distribution patterns across 16 metagenomic samples. Finally, we visualized all this information using the program `anvi-interactive`. User tutorials for the meta-pangenomic workflow is available online at the URL http://merenlab.org.

